# Optical constraints on two-photon voltage imaging

**DOI:** 10.1101/2023.11.18.567441

**Authors:** F. Phil Brooks, Hunter C. Davis, J. David Wong-Campos, Adam E. Cohen

## Abstract

**Significance:** Genetically encoded voltage indicators (GEVIs) are a valuable tool for studying neural circuits *in vivo*, but the relative merits and limitations of one-photon (1P) vs. two-photon (2P) voltage imaging are not well characterized.

**Aim:** We consider the optical and biophysical constraints particular to 1P and 2P voltage imaging and compare the imaging properties of commonly used GEVIs under 1P and 2P excitation.

**Approach:** We measure brightness and voltage sensitivity of voltage indicators from commonly used classes under 1P and 2P illumination. We also measure the decrease in fluorescence as a function of depth in mouse brain. We develop a simple model of the number of measurable cells as a function of reporter properties, imaging parameters, and desired signal-to-noise ratio (SNR). We then discuss how the performance of voltage imaging would be affected by sensor improvements and by recently introduced advanced imaging modalities.

**Results:** Compared to 1P excitation, 2P excitation requires ∼10^4^-fold more illumination power per cell to produce similar photon count rates. For voltage imaging with JEDI-2P in mouse cortex with a target SNR of 10 (spike height:baseline shot noise), a measurement bandwidth of 1 kHz, a thermally limited laser power of 200 mW, and an imaging depth of > 300 μm, 2P voltage imaging using an 80 MHz source can record from no more 12 cells simultaneously.

**Conclusions:** Due to the stringent photon-count requirements of voltage imaging and the modest voltage sensitivity of existing reporters, 2P voltage imaging *in vivo* faces a stringent tradeoff between shot noise and tissue photodamage. 2P imaging of hundreds of neurons with high SNR at depth > 300 μm will require either major improvements in 2P GEVIs or qualitatively new approaches to imaging.

## Introduction

A longstanding dream in neuroscience has been to record the membrane potential of hundreds or thousands of neurons simultaneously in a behaving animal. Such measurements could reveal functional connections, probe input-output properties of cells and of microcircuits, and help discern principles of neural computation. Recent advances in GEVIs have substantially improved their signal-to-noise ratio (SNR), enabling recordings from dozens of cells in superficial tissue using one-photon imaging^1–3^. There have also been improvements in instrumentation^4,5^ and reporters^6^ for 2P voltage imaging, enabling voltage imaging at depths up to 500 μm, though the number of simultaneously recorded cells at this depth remains < 3. Most applications of voltage imaging *in vivo* have been with 1P excitation ^7–10^, whereas for Ca^2+^ imaging, 2P microscopy is the dominant approach.^11^ This raises the question: what are the scaling properties and relative merits of 1P vs 2P voltage imaging *in vivo*? How can a researcher considering a voltage imaging experiment decide which approach to use?

The physical requirements of Ca^2+^imaging and voltage imaging differ substantially, so intuitions may not transfer. For Ca^2+^ imaging, typical events last 100 – 500 ms and have amplitudes of ΔF/F ∼100%. Signals come from the bulk cytoplasm. For voltage imaging, action potentials last ∼0.3 – 2 ms, and typically have amplitudes of ΔF/F ∼ 10 - 30%, though subthreshold events can be 100-fold smaller. Signals are localized to the cell membrane. Thus, the key challenge in voltage imaging is to acquire adequate SNR and imaging speed in the presence of shot-noise and motion artifacts, while maintaining tissue-safe laser powers.

Here we explore, with the support of modeling and data from representative voltage indicators, how molecular and optical parameters affect the balance between SNR, number of simultaneously recorded cells, and tissue damage. Many of the arguments about scaling of noise^12,13^ and 2P signal^14,15^ are found elsewhere in the literature, but with the recent publications seeking to reach^5,6,16–22^ or transcend^23–27^ these limits, we believe a consolidation of the arguments with a specific application to voltage imaging is warranted.

Our results support the preference for 1P over 2P imaging at shallow depths and the use of 2P voltage imaging at depths where 1P recordings are inaccessible due to light scattering. However, at depths beyond the 1P limit, 2P voltage imaging signals are severely constrained by thermal and shot-noise limits. We address the potential of advanced instrumentation and analysis techniques to improve performance beyond current limits.

## Materials and Methods

### HEK cell culture

HEK293T cells were maintained at 37 °C, 5% CO2 in Dulbecco’s Modified Eagle Medium (DMEM) supplemented with 10% fetal bovine serum, 1% GlutaMax-I, penicillin (100 U/mL), streptomycin (100 mg/mL). For maintaining or expanding the cell culture, we used 35 mm TC-treated culture dish (Corning). For each imaging experiment, cells in one 35 mm dish were transiently transfected with the construct to be imaged using PEI in a 3:1 PEI:DNA mass ratio. For all the imaging experiments, cells were replated on glass-bottomed dishes (Cellvis, D35-14-1.5-N) 36 hours after transfection. Imaging was performed ∼6 hours after replating. Before optical stimulation and imaging, the medium was replaced with extracellular (XC) buffer containing 125 mM NaCl, 2.5 mM KCl, 3 mM CaCl2, 1 mM MgCl2, 15 mM HEPES, 30 mM glucose (pH 7.3). For BeRST1 experiments in HEK cells, cells were stained with 1 μM BeRST1 for 30 min prior to 3x wash with XC buffer before imaging.

### Microscope and illumination control

All imaging was performed using Luminos bi-directional microscopy control software^28^ on a custom-built upright microscope equipped with 1P and 2P illumination paths and a shared emission path to an sCMOS camera (Hamamatsu, ORCA-Flash 4.0 v2). The 1P illumination path contained a 488 nm laser (Coherent OBIS), a 532 nm laser (Laserglow LLS-0532) and a 635 nm laser (Coherent OBIS). The outputs of the 488 nm and 532 nm lasers were modulated using a multichannel acousto-optic tunable filter (Gooch & Housego PCAOM NI VIS driven by G&H MSD040-150). The 635 nm laser was modulated using its analog modulation input and an external neutral density filter wheel (Thorlabs). The 488 nm and 532 nm lasers were patterned via a Digital Micromirror Display (Vialux V-7001 V-module). To convert illumination intensity to power per cell, we approximated HEK cells as circles with a 10 μm diameter, e.g., an intensity of 1 W/cm^2^ corresponded to 0.8 μW per cell.

The two-photon illumination path comprised an 80 MHz tunable femtosecond laser (InSight DeepSee, Spectra Physics), an electro-optic modulator (ConOptics 350-80-02) and two galvo mirrors for steering (Cambridge Technology 6215H driven by 6671HP driver). Power calibration was performed with a Thorlabs PM400 power meter with a photodiode-based sensor (S170C) and a thermal sensor (S175C) for 1P and 2P illumination, respectively. Electrical stimuli and measurements were performed using a National Instruments 6063 PCIe DAQ. All imaging was performed with a 25x water immersion objective (Olympus XLPLN25XWMP2, 2mm working distance, NA 1.05). At each wavelength, the dispersion was adjusted to maximize the 2P fluorescence signal. The wavelengths for 2P excitation of QuasAr6a and BeRST1 were chosen to drive the S_0_ to S_2_ transition rather than the S_0_ to S_1_ transition because the S_2_ transition was stronger.

### Measuring voltage-sensitive fluorescence in vitro

All imaging and electrophysiology experiments were performed in XC buffer. Concurrent whole-cell patch clamp and fluorescence recordings were acquired on the microscope described above. Filamented glass micropipettes were pulled to a tip resistance of 5–8 MΩ and filled with internal solution containing 125 mM potassium gluconate, 8 mM NaCl, 0.6 mM MgCl_2_, 0.1 mM CaCl_2_, 1 mM EGTA, 10 mM HEPES, 4 mM Mg-ATP and 0.4 mM Na-GTP (pH 7.3), adjusted to 295 mOsm with sucrose. Whole-cell patch clamp recordings were performed with an Axopatch 200B amplifier (Molecular Devices). Fluorescence was recorded in response to a square wave from - 70 mV to +30 mV.

### In-house AAV packaging

AAV2/9 JEDI-2P vectors were packaged in house based on a published protocol.^29^ Briefly, 50∼70% confluent HEK293T cells grown in DMEM supplemented with 5% FBS were triple transfected with pHelper, pAAV ITRexpression, and pAAV Rep-Cap plasmids using acidified (pH 4) PEI (DNA:PEI ratio 1:3) in 1∼2 T175 flasks (∼2 × 10^7^ cells/flask). The AAV-containing medium was harvested on Day 3, and the AAV-containing medium and cells were harvested on Day 5. For the second harvest, AAVs were released from the cells with citrate buffer (55 mM citric acid, 55 mM sodium citrate, 800 mM NaCl, 3 mL per flask). The two harvests were then combined and precipitated with PEG/NaCl (5x, 40% PEG 8000 (w/v), 2.5 M NaCl, 4 °C overnight). The low-titer virus was then purified with chloroform extraction (viral suspension and chloroform 1:1 (v/v)), aqueous two-phase partitioning (per 1 g of the AAV-containing supernatant, add 5 g of 20% (NH_4_)_2_SO_4_ solution and 1.5 g of 50% PEG 8000 solution, and iodixanol discontinuous gradient centrifugation (15%, 25%, 40%, and 54% iodixanol gradient prepared from OptiPrep (60% (w/v) Iodixanol, Axis-Shield PoC AS). The purified AAV titer was determined via qPCR (SYBR Green, primer for forward ITR: 5’-GGAACCCCTAGTGATGGAGTT-3’; primer for reverse ITR sequence 5’-CGGCCTCAGTGAGCGA-3’).

### In vivo imaging

All animal experiments were approved by the Institutional Animal Care and Use Committee of Harvard University. The cranial window surgery for *in vivo* imaging was based on previously published protocols^7^. Briefly, an adult CD1 mouse was injected with 50 nL viral mix in 4 sites in the whisker barrel cortex at 100, 200, 300, 400, and 500 μm below the tissue surface. Viral mix had a final concentration of 5e12 vg/mL pAAV-EF1a-DIO-JEDI-2P-Kv2.1motif and 1e11 CamKII-Cre in AAV 2/9. A cranial window and mounting plate were then installed over the injection sites. Two weeks after surgery and injection, a head fixed CD1 mouse was imaged at 1.5% isoflurane with the dose adjusted to maintain a stable breathing rate. The mouse was kept on a heating pad (WPI ATC2000) to maintain stable body temperature at 37 °C and its eyes were kept moist using ophthalmic eye ointment. 2P imaging was performed at a wavelength of 930 nm. The mouse was imaged for 2 hours after which it recovered in less than 10 minutes.

Light for 1P imaging *in vivo* was patterned via a digital micromirror device to selectively illuminate the targeted cell body, as described above.

### Data analysis

All analysis was performed in Matlab. Cell fluorescence was measured from regions of interest (ROIs) manually selected to lie on the cell membrane. 1P and 2P recordings from the same cell used identical analysis ROIs. Fluorescence from cell-adjacent illuminated areas was used to estimate background.

To estimate shot noise for an ideal detector, we converted camera counts to incident photons by dividing by the camera quantum efficiency (QE = ∼67% @ 525 nm) and multiplying by the conversion factor (CF = 0.46 photoelectrons/digital count).

## Results

### Shot noise constrains functional fluorescence imaging

Shot noise imposes a fundamental limit on imaging performance. A source that generates, on average, *N* detected photons, will have fluctuations with standard deviation 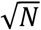. To detect a 1 ms spike (ΔF/F ∼ 10%) with an SNR of 10, requires determining fluorescence to 1% precision in 1 ms.

We adopt 1% precision in 1 ms as a reasonable standard for a high-SNR voltage recording. Due to shot noise, this standard requires detecting at least 10^4^ photons/ms, or 10^7^ photons/s.

More generally, imaging an event of magnitude ΔF/F = β with a given SNR requires determining fluorescence to a precision of β/SNR. The photon flux (Г) required for a measurement rate (*f*), signal level (β), and SNR is:

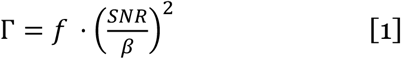

If one wishes only to detect spikes in pyramidal cells, one might tolerate SNR = 3 and *f* = 400 Hz. A highly sensitive GEVI could give β = 0.2 for a spike.^6^ Under these conditions, the minimum detection rate is *Г* = 9×10^4^ photons/s. In comparison, for a typical calcium imaging scenario, SNR = 10, β = 1 and *f* = 30 Hz,^30^ implying *Г* = 3000 photons/s, 30-fold less than even low-SNR spike detection via voltage imaging. Thus, the brief duration of voltage spikes and the low fractional sensitivity of existing voltage indicators conspire to make voltage imaging a very photon-greedy technique.

Filtering in space or time can increase the effective value of *N* at a given pixel, at the cost of lower spatial or temporal resolution, but filtering does not change the 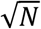 shot noise scaling. We discuss advanced analysis techniques below.

### 1P vs 2P excitation of commonly used voltage indicators

We compared the brightness of commonly used voltage indicators in HEK293T (HEK) cells under alternating wide-field 1P illumination and 2P spiral illumination with an 80 MHz pulsed laser, using a shared detection path for both modalities to ensure equal photon detection efficiencies (Fig. 1a-c; Methods). We compared: a voltage sensitive dye (BeRST1^31^), an opsin-derived GEVI imaged via intrinsic retinal fluorescence (QuasAr6a^32^), a chemigenetic FRET-opsin GEVI (Voltron2^33^), and two GEVIs that couple a voltage sensitive phosphatase (VSP) to a circularly permuted partner fluorophore (ASAP3^4^ and JEDI-2P^6^).

**Figure 1.**
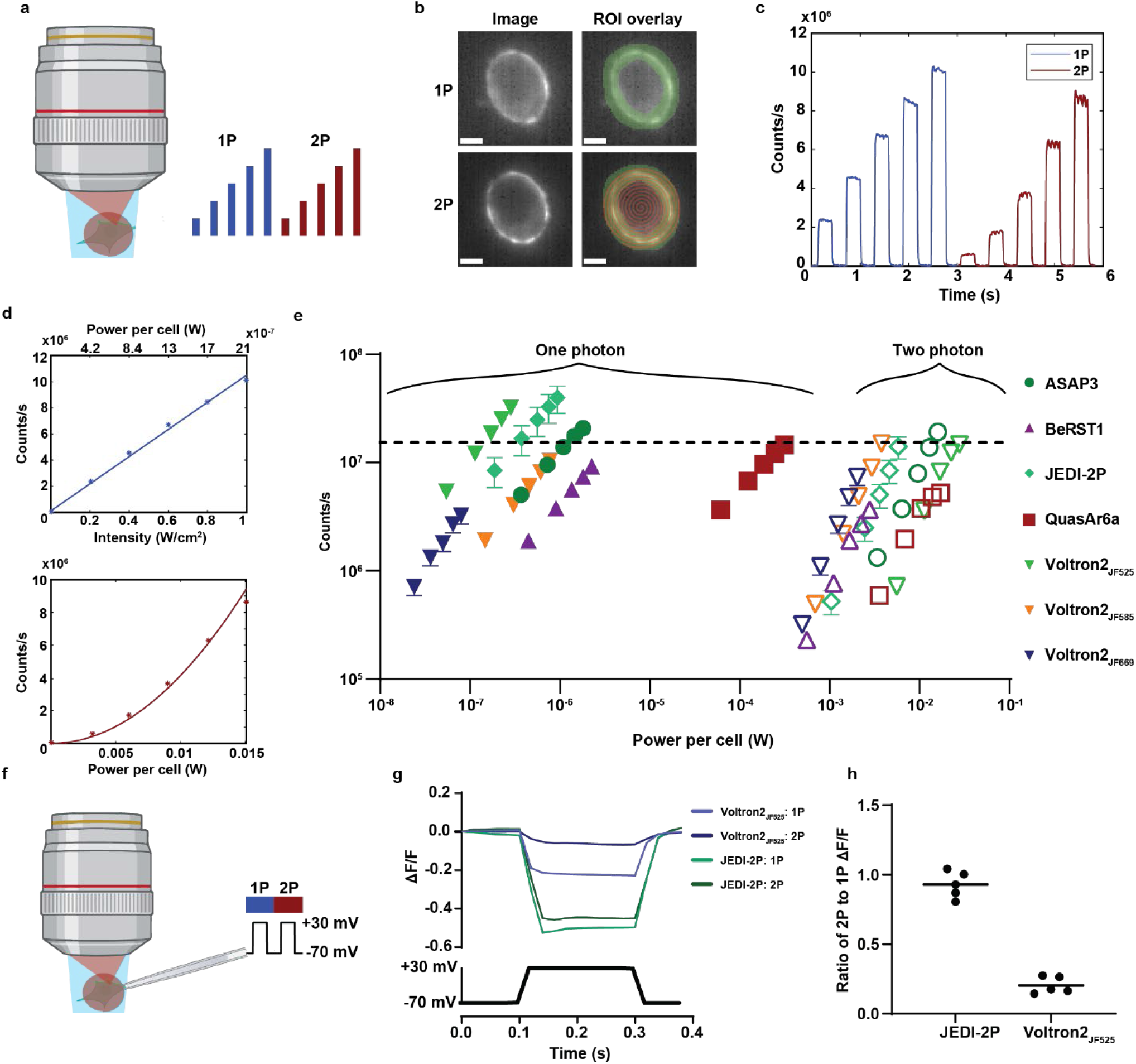
Comparison of 1P and 2P brightness and sensitivity of fluorescent voltage indicators. A) Diagram of experiment. HEK cells were sequentially illuminated with wide-field 1P light in steps of increasing intensity, then by spiral scan 2P steps of increasing intensity. B) Example HEK cell expressing the GEVI ASAP3. 2P spiral scan pattern shown in red and analysis ROI shown in green. Scale bars 5 μm. C) Example single-trial data for cell expressing ASAP3. D) Top: fluorescence in c as a function of 1P intensity with linear fit. Bottom: fluorescence as a function of 2P power with quadratic fit. E) Log-log plot of count rate vs. optical power on the cell for seven voltage indicators. 1P data: filled symbols, 2P data: empty symbols. Error bars are SEM from at least *n* = 8 cells. The excitation wavelengths used for 1P (2P) excitation of each of the reporters were: ASAP3 488 nm (930 nm), BeRST1 635 nm (850 nm), JEDI-2P 488 nm (930 nm), QuasAr6a 635 nm (900 nm), Voltron2_525_ 488 nm (930 nm), Voltron2_585_ 594 nm (1100 nm), Voltron2_669_ 635 nm (1220 nm). A horizontal line is shown at 1.5×10^7^ counts/s, equivalent on our camera to 10^7^ impinging photons/s. F) Whole-cell patch clamp protocol for measuring voltage sensitivity under 1P and 2P excitation. G) Average voltage responses of JEDI-2P and Voltron2_525_ under 1P and 2P illumination. H) Ratio of voltage contrast under 2P vs. 1P illumination for JEDI-2P and Voltron2_525_, *n* = 5 cells per construct.

As expected, the fluorescence scaled linearly with 1P illumination intensity and quadratically with 2P intensity (Fig. 1d). For all but one indicator, 2P illumination required at least 10^4^-fold greater time-averaged power per cell to achieve comparable counts to 1P illumination (Fig. 1e). The ratio of 2P to 1P powers for QuasAr6a was only ∼300, due to the requirement for high intensity 1P illumination and selection rules which favor 2P over 1P excitation in opsins.^34^ Consistent with prior reports,^1^ we found that chemigenetic indicators were brighter under both 1P and 2P illumination than their purely protein-based counterparts, though our measurements did not assign the relative contributions of expression level vs. per-molecule brightness.

The huge difference in optical power requirement between 1P and 2P (80 MHz) excitation is consistent with published reports: 2P imaging of JEDI-2P was reported at a power of 9 – 12 mW/cell^6^, whereas 1P imaging of similar GFP-based GEVIs is typically performed at 1 – 10 W/cm^2^,^4,6^ corresponding to 1 – 10 μW/cell. Estimates based on tabulated 1P and 2P absorption coefficients^35^ give a similar factor of ∼10^4^ difference in power efficiency (Supplementary text).

We then measured voltage sensitivity of one representative from each GEVI family, comparing 1P and 2P illumination. Using whole-cell voltage clamp in HEK cells (Fig. 1f), we found that the contrast (ΔF/F per 100 mV) of the VSP-based JEDI-2P was similar for 1P and 2P illumination. The opsin-based chemigenetic indicator Voltron2^525^ showed voltage sensitivity under 1P but not 2P illumination (Fig. 1g-h). Loss of voltage sensitivity under 2P illumination of FRET-Opsin GEVIs has been reported previously^36^. We recently determined the photophysical mechanism underlying this effect,^37^ but did not pursue 2P imaging of FRET-Opsin GEVIs here.

### Testing the dependence of 1P and 2P signals as a function of depth

To characterize the depth dependence of 1P and 2P voltage imaging in brain tissue, we sparsely expressed soma-localized JEDI-2P in mouse cortex. Chien *et al*. previously showed that for voltage imaging in brain tissue, restricting 1P illumination to the soma led to an ∼8-fold improvement in signal-to-background ratio compared to wide-field illumination, by minimizing background from off-target illumination.^38^ We thus compared soma-targeted 1P imaging and raster-scanned 2P imaging at several depths (Fig. 2a-b; Methods).

**Figure 2.**
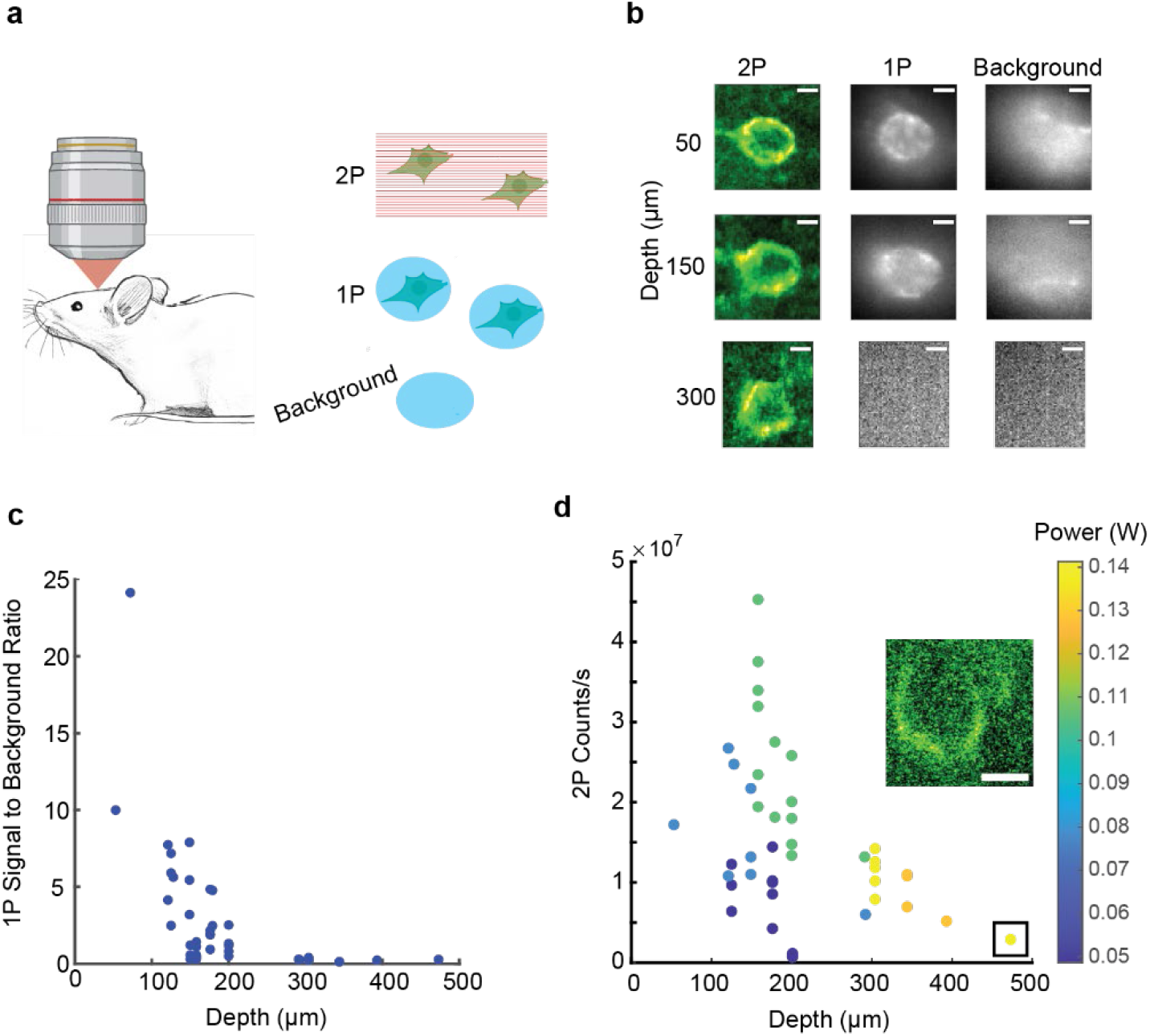
Depth-dependent 1P and 2P signal in brain. A) Experimental protocol. In a mouse expressing JEDI-2P, raster-scanned 2P (*λ* = 920 nm) and DMD-patterned 1P imaging (*λ* = 488 nm) were alternately applied to neurons at different depths. 1P illumination patterned to cell-free regions was used to estimate background signal. B) Example 1P and 2P images of cells at three depths. Scale bars 5 μm. C) Estimated signal-to-background ratio for *n* = 43 neurons under patterned 1P illumination. D) Mean count rate from the membranes of *n* = 43 neurons under 2P illumination. Color indicates excitation power. Inset shows cell at 473 μm depth (boxed on the graph). Scale bar 5 μm.

With 1P illumination, JEDI-2P-expressing cells were resolvable down to *d* ∼ 200 μm. The challenge for 1P imaging at greater depths was not shot noise, but rather decrease in signal-to-background ratio. At depths > 200 μm, cells were not distinguishable from background by 1P imaging with patterned 488 nm excitation (Fig. 2c).

With 2P illumination, cells were resolvable to *d* = 473 μm, the greatest depth we tested (Fig. 2b). As the cell depth increased, we increased the 2P laser power to maintain sufficient count rate to resolve the cells, up to *P*_0_ = 140 mW at *d* = 473 μm.

### Thermal limits to 2P excitation power in brain tissue

The heating caused by 2P illumination can transiently perturb neural function, and at high levels can damage tissue. Most ion channel gating properties have a Q_10_ (i.e. ratio of rates at temperatures separated by 10 °C) between 1 and 3.^39^ Changes of 1°C can cause changes in neuronal firing rates.^40^ In rodents, brain temperature may fluctuate under physiological conditions by up to 4 °C.^41^ Podgorski and Ranganathan^42^ found lasting damage after continuous illumination of a 1 mm^2^ scan at 250 mW, corresponding to a steady-state temperature change of ∼5 °C.

The relation between laser power and heating depends on scan area, scan pattern, and measurement duty cycle. For a 1 mm^2^ square scan pattern, Podgorski and Ranganathan found steady-state temperature coefficients between 0.012 and 0.02 °C/mW at wavelengths from 800 – 1040 nm, equivalent to a temperature rise of < 2 °C at 100 mW illumination. They simulated the dependence of temperature rise on scan area and found a weak dependence. Their results predict a maximum temperature coefficient of 0.03 °C/mW for a square scan of side length 20 μm (representing a single neuronal soma). The precise value of this coefficient depends both on the brain region and the wavelength^43^. Finally, Podgorski and Ranganathan show that reducing the illumination duty-cycle to 10 s on, 20 s off, allowed brighter illumination to be used during “on” periods, while still keeping time-averaged heating beneath the damage threshold. Some experiments may permit low duty-cycle imaging while others may not. Hereafter we use 200 mW as a reasonable upper bound on the power, acknowledging that this limit may be higher (or lower) depending on many experimental details.

### Estimating measurable cells as a function of depth

Shot noise places an upper bound, 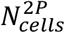, on the number of cells which can be measured simultaneously via 2P illumination with a given illumination power and SNR. Under a protocol which sequentially visits single cells, 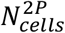 depends on both the brightness and voltage sensitivity (see Supplementary Information for derivation):

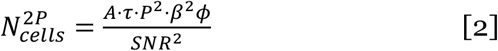

Here *A* is the brightness coefficient derived from the HEK cell experiments (Eq. S1), *τ* is the integration time, *P* is the laser power at the focus, *β* = ΔF/F per spike, *ϕ* is the fraction of the scan that intersects with the cell membrane (i.e. the imaging duty cycle), and *SNR* is the target ratio of the spike amplitude to shot noise. The value of *A* is specific to the laser repetition rate, pulse width, focal parameters, and detection optics. We discuss the effects of varying these parameters below. For an analysis that includes the effect of light scattering on depth-dependent collection efficiency, see ^15^.

The parameter *ϕ* approaches 1 for perfectly membrane-targeted illumination. To estimate *ϕ* for a raster scan over a bounding box around a single cell body, we examined 2P images of pyramidal cells with membrane-targeted fluorescent tags in cortex layer 2/3. In these images, the membrane-labeled area fraction within the bounding box was *ϕ*_*bb*_ = 0.18 ± 0.07 (mean ± std, *n* = 10 cells). For a raster scan over multiple sparsely expressing cells, *ϕ* is lower than *ϕ*_*bb*_ by a factor of the sparsity. The low values of *ϕ* for raster-scanned 2P imaging are a consequence of the membrane-localized signal and highlight the importance of membrane-targeted illumination. However, precise targeting of the illumination to the membrane increases sensitivity to motionM artifacts. The ULoVE technique brackets the membrane with pairs of spots and thereby mitigates the effect of small motions, at the cost of less-than-optimal membrane targeting of the spots.^4^

Eq. 2 can be used to predict the scaling of 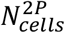 as a function of depth, *d*, for 2P voltage imaging. The power at a laser focus decays exponentially with *d*, with an extinction length, *l*_e,_ in brain tissue. At *λ* = 920 nm *l*_e_ is between 112^15,44^ and 155 μm.^45^ Due to the quadratic dependence of 2P signal on focal intensity, the 2P signal decays with a length constant of *l*_e_/2. Substituting 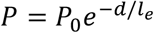 into Eq. 2 implies a decay in 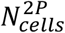 by a factor of 10 for every 130 - 180 μm increase in *d* (Fig. 3a), assuming constant power *P*_0_ into the tissue.

**Figure 3.**
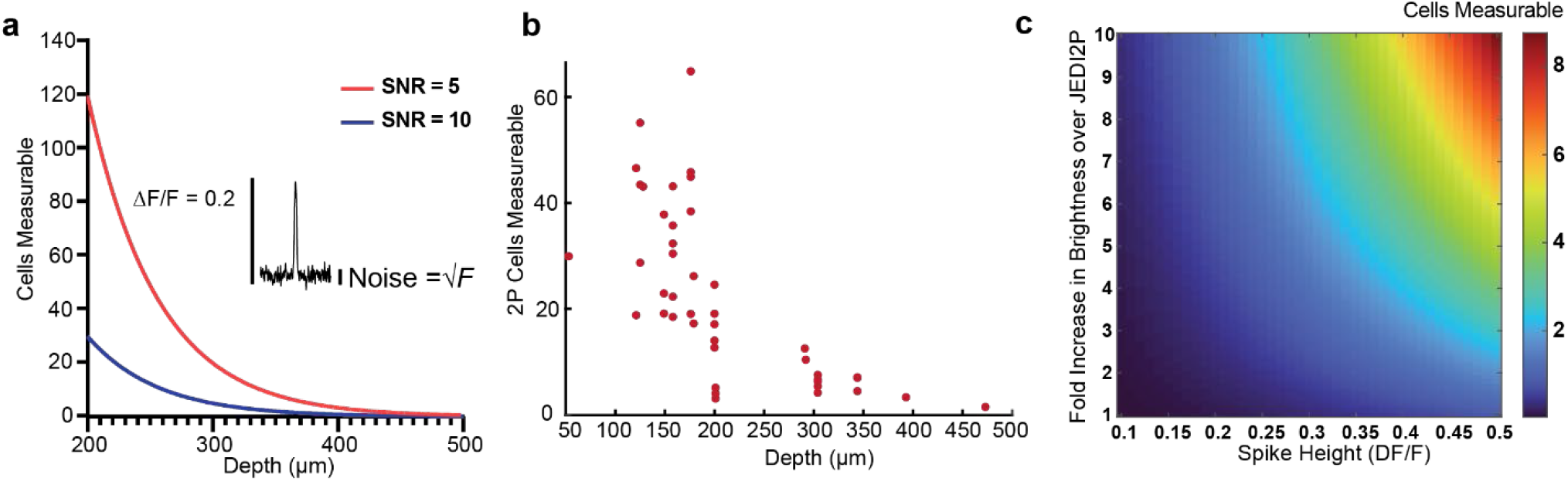
Scaling of 2P voltage measurements with depth and with GEVI properties. A) Predicted number of simultaneously measurable cells as a function of depth, based on brightness derived from HEK cells expressing JEDI-2P (Fig. 1). We assumed a spike contrast of *β* = 0.2, target SNR of 5 (blue) or 10 (red) in an integration time *τ* = 1 ms, a total power *P*_*0*_ = 200 mW, targeting fraction *ϕ* = 1, scattering length *l*_e_ = 112 μm^44^, and a detector with perfect quantum efficiency. B) Predicted number of simultaneously measurable cells at SNR = 10 and 200 mW power for each experimentally measured single-cell count-rate reported in (Fig. 2d). C) Number of simultaneously measurable cells under 2P illumination at a depth of 500 μm, assuming brightness and contrast improvements of future GEVIs, target SNR of 10, and all other parameters as in (A).

We applied Eq. 2 to the instantaneous count rates measured from the cell membranes (Fig. 2d) to estimate 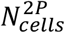 for *ϕ* ∼ 1, i.e., a perfectly membrane-targeted annular scan pattern. We converted the digital camera counts to collected photon rates to provide an upper performance bound for a perfect detector. Modern back-illuminated sCMOS cameras have detection efficiencies that approach 100%. We assumed *β* = 0.2, based on the reported spike response of JEDI-2P^6^ and a target shot noise-limited SNR of 10 in a 1 kHz bandwidth, and extrapolated the count rates to an input power of *P*_0_ = 200 mW. The estimated number of measurable cells decreased quickly at *d* > 200 μm and dropped below three at *d* > 470 μm and input power 200 mW (Fig. 3b). These results are similar to the prediction from the simple model using the 2P count rates from our HEK cell experiments (Fig. 3a, blue line).

The palette of available voltage indicators is continually improving.^46^ We therefore considered the scaling of 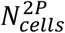 with changes of brightness and spike height (*β*).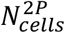 depends linearly on *A* and quadratically on *β* (Fig. 3c). An order-of-magnitude increase in molecular brightness coupled with a 2.5 fold increase in *β* compared to JEDI-2P could enable high SNR measurement of >8 cells at depths up to 500 μm using the optical configuration we considered above.

### Effect of GEVI kinetics

Some GEVIs have response times that are slow compared to the duration of a spike. On one hand, this blunts the amplitude of the spike response; on the other, it permits one to average for longer to detect whether a spike has occurred (assuming that the interval between spikes remains long compared to the recovery time of the GEVI). Here we analyze this tradeoff.

Consider a GEVI subjected to a voltage step which induces a steady-state change in fluorescence, 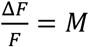. Assume that the GEVI responds to a voltage step of duration *t* with exponential response time constants *τ*_*on*_ and *τ*_*off*_ (Fig. 4a,b). We can then write the total integrated fractional response, *R*, as (See Supplement for derivation):

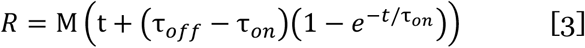

**Figure 4.**
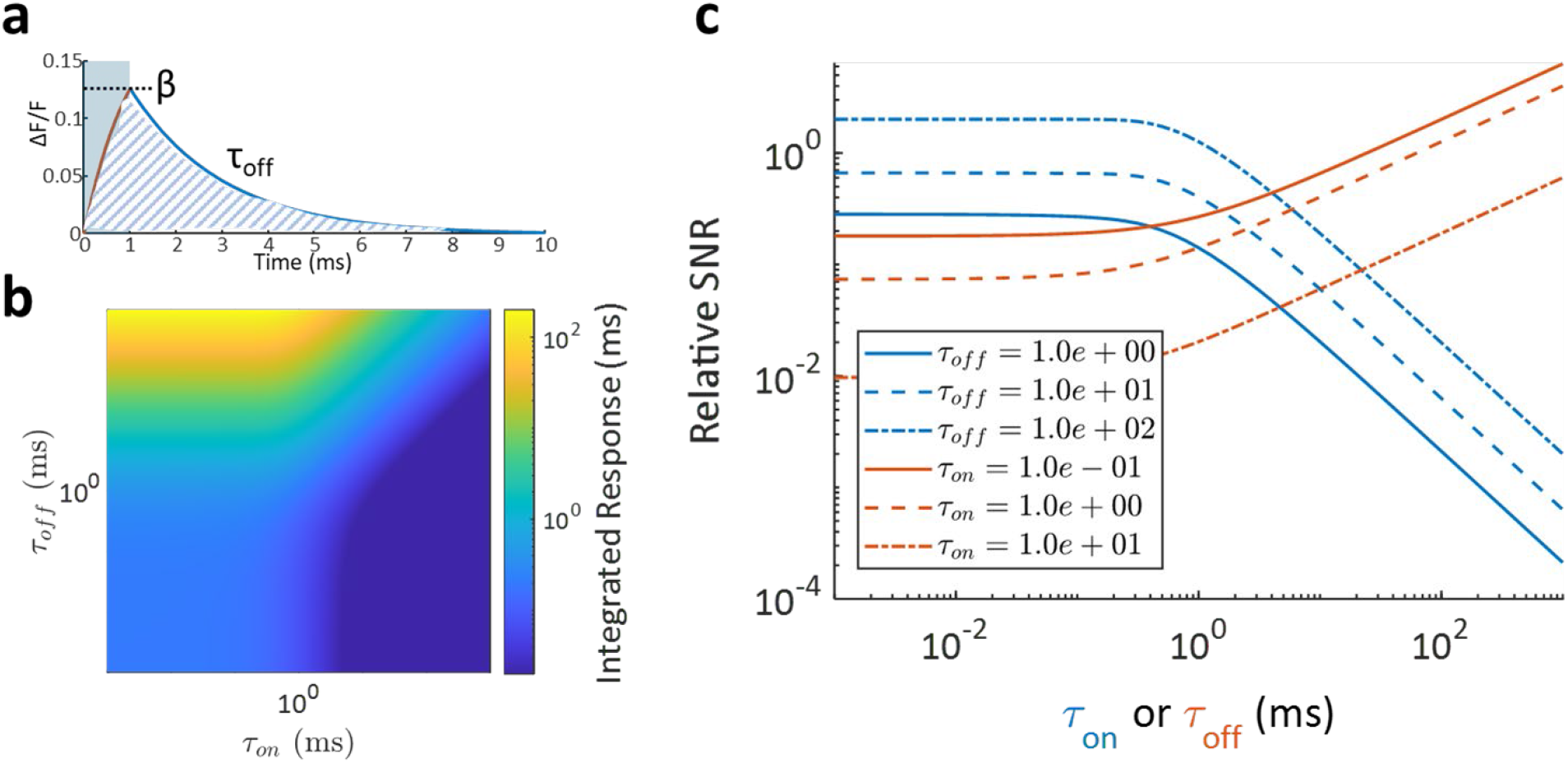
Effect of reporter kinetics on signal. A) Response of a reporter to a 1 ms voltage pulse. An exponential rise with time constant *τ*_*on*_ reaches a maximum of *β*, followed by an exponential decay with time constant *τ*_*off*_. The signal comprises the total area under both response phases (cross-hatched). B) Area under the curve in (A) as a function of *τ*_*on*_ and *τ*_*off*_, keeping constant pulse duration (1 ms) and steady-state voltage sensitivity (*β*_*ss*_ = 0.2). Increasing *τ*_*off*_ allows longer integration time, while increasing *τ*_*on*_ truncates the response. C) Effects on SNR of changing *τ*_*on*_ or *τ*_*off*_ while keeping the other fixed. The fixed parameter is shown in the legend and the variable one is indicated by the x-axis. At large *τ*_*on*_, SNR ∼ 1/*τ*_*on*_. At large *τ*_*off*_, SNR ∼ *τ*_*off*_ ^½^.

Eq. 3 assumes collection of the entire tail of the decay and thus provides an upper bound to the signal. For a finite integration time equal to t + τ_*off*_, the relation of SNR and *R* is (See Supplement for derivation):

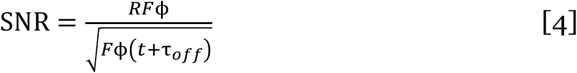

A large *τ*_*on*_ will decrease the magnitude of the response β to a short electrical spike, decreasing instantaneous SNR proportionally. A large *τ*_*off*_, however, increases the duration of the impulse response, increasing the duration of the signal that can be integrated and leading to an increase in SNR proportional to *τ*_*off*_ ^½^ (Eq. S7, Fig. 4c). When *τ*_*on*_ and *τ*_*off*_ can be independently chosen, *τ*_*on*_ should be minimized (with diminished effect once *τ*_*on*_ is below the spike width) and *τ*_*off*_ maximized (while remaining short compared to the inter spike interval). Often, these time constants are biophysically related. If *τ*_*on*_ ≈ *τ*_*off*_ = *τ*, then *SNR* ∼*τ*^−½^ (see Eq. S7). That is, faster GEVIs are better than slower ones in terms of shot noise-limited SNR, all else being equal.

### Effect of optical parameters on 2P fluorescence

Advances in 2P voltage imaging typically have two aims: 1) increasing the number of cells, *N*, which are sampled with high enough revisit rate to capture all spikes; and 2) increasing fluorescence per cell to improve the shot noise-limited SNR. In many cases, these two aims are in tension.

#### Changing numerical aperture

We distinguish between numerical aperture of excitation (NA_e_) and of collection (NA_c_). Often, NA_c_ is set by the objective NA, while NA_e_ may be lower as a result of underfilling the objective back aperture. The photon detection efficiency, *PDE*, scales as *PDE* ∼ *NA*_*c*_^2^. The effect of excitation numerical aperture (NA_e_) on the 2P signal depends on the sample geometry. Within the Gaussian beam approximation (Fig. 5a), the width of the focus scales as

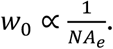

**Figure 5.**
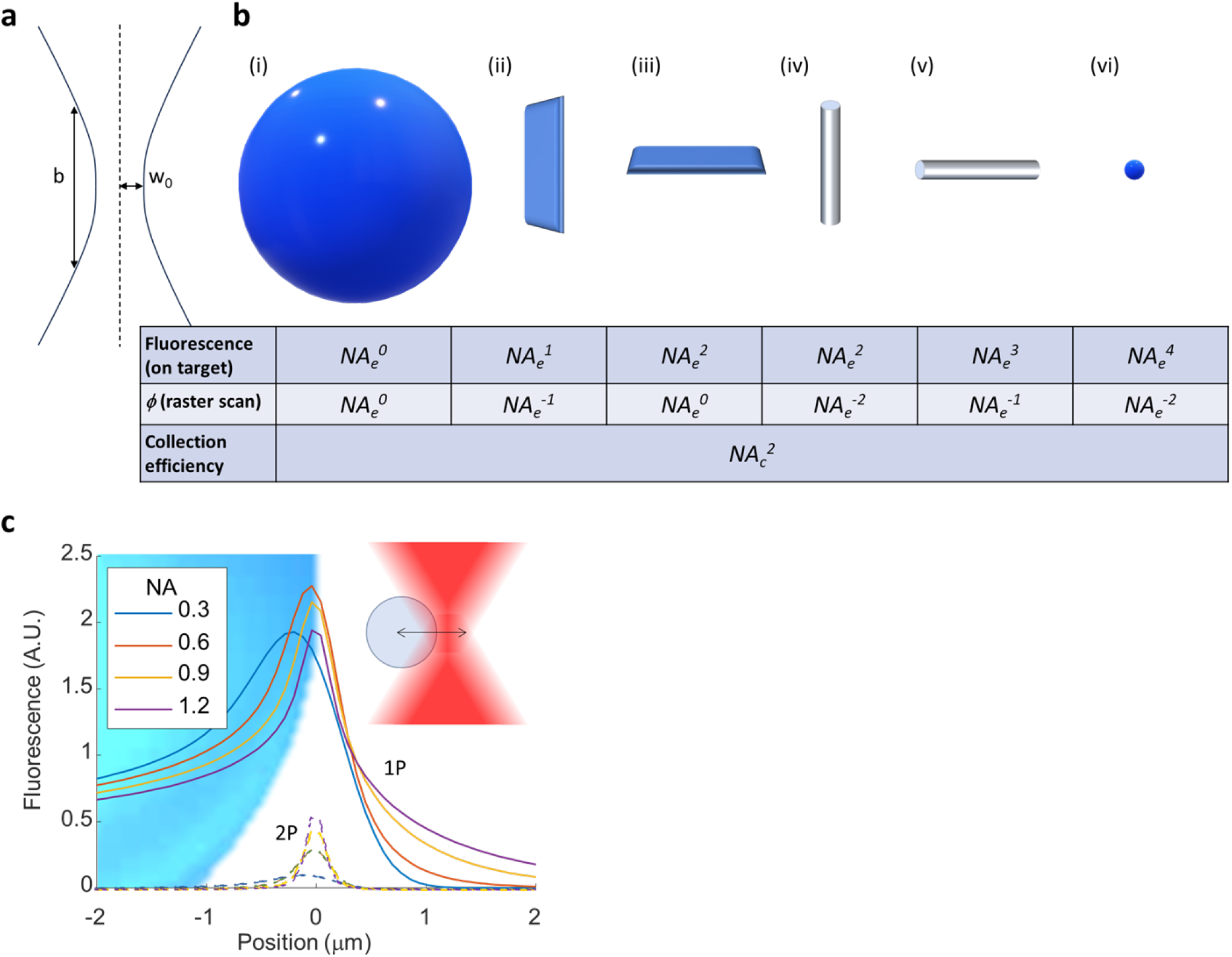
Scaling of fluorescence with numerical aperture, sample geometry, and excitation modality. A) Geometry of a Gaussian beam, showing the width (w_0_ ∼ 1/*NA*) and waist (*b* ∼ 1/*NA*^2^). B) Scaling of total 2P fluorescence as a function of excitation NA_e_ for different sample geometries. All slender dimensions are assumed to be << *w*_0_ and all extended dimensions are assumed to be >> *b*. In a planar raster scan, the fraction of time that a sub-wavelength structure is excited, *ϕ*, depends on the focus width and hence the NA_e_. In cases (iii), (v), and (vi) we assume that the object is perfectly in focus, i.e., in the axial plane where focus size is minimum. The collection efficiency for all geometries depends on the collection solid angle ∼*NA*^2^_c_. To calculate total signal for targeted illumination, multiply the first and third lines; for a raster scan, multiply all three lines. C) Total fluorescence evoked by the intersection of a laser focus and a spherical membrane, 10 μm diameter. We compared 1P and 2P excitation with equal NA_c_ and NA_e_, with powers adjusted to match per-molecule excitation rates at the focus at NA = 1.2. The much smaller 2P focal volume led to 5.3-fold smaller maximum fluorescence at the highest NA (and even greater discrepancy at lower NA) and 3-fold greater sensitivity to misalignment, compared to 1P excitation.

The intensity, *I*, at the focus scales inversely with the cross-sectional area 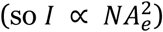, and the rate of 2P excitation per molecule, *E*, scales with the intensity squared. Hence

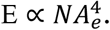

The axial extent of the beam waist scales as

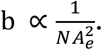

The total signal from a volume element depends on the number of fluorophores excited. In a three-dimensional bulk solution — an approximation of 2P Ca^2+^ imaging when the beam waist is significantly smaller than a single cell — the volume scales approximately as 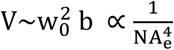. The total fluorescence emission Г_2P_ scales as *V*×*E*. Hence, in bulk solution, 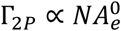, so that the total collected fluorescence, *F*, scales only with NA_c_ as 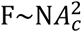.

In contrast, for 2P voltage imaging, the fluorescence rate from the sample scales with NA_e_ raised to a power between 1 and 3, depending upon the orientation and geometry of the membranes in the focal spot (Fig 5b, first row). This scaling argues strongly for maximizing the NA_e_ for 2P voltage imaging. On the other hand, a smaller excitation spot leads to (a) a higher rate of photobleaching and (b) greater sensitivity to misalignment between the focus and the sample, e.g., from sample motion (Fig. 5c).

#### Advanced scanning modalities

State-of-the-art techniques for high-speed 2P microscopy often involve shaping and splitting the excitation in space and/or time^4,5,20,47,48^. The effect of pulse splitting depends on whether the illumination is limited by total power into the sample or by nonlinear photodamage at the laser focus. In the case of limited total power, a diffraction-limited focus provides the highest possible 2P excitation rate and hence SNR. In the case of limited focal energy density, pulse splitting may be advantageous. Here we discuss the effects on optical SNR of changes to instrumentation.

#### Spatial and temporal focal multiplexing

Some 2P imaging modalities use beam splitters to split each laser shot into a series of pulses which arrive sequentially at different locations in the sample.^17,20,23,47^ Other modalities expand the laser beam to cover more than a single diffraction-limited spot (e.g. multifocal^4,16,23,48,49^, temporal focusing^5,48,50^, or Bessel beam^51^ microscopies). Consider a laser with repetition rate *f* that is split into *N* diffraction-limited spots (either temporal or spatial splitting). We define the effective repetition rate, *f*_eff_ = *N*×*f*. The number of diffraction-limited spots imaged per second is proportional to *N*, but the excitation rate per spot is proportional to 1/*N*^2^, assuming constant total power into the sample. Hence, the fluorescence count rate scales as 1/*N* (Fig. 6a), and the number of measurable cells at fixed SNR and input power scales as 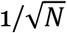.

**Figure 6.**
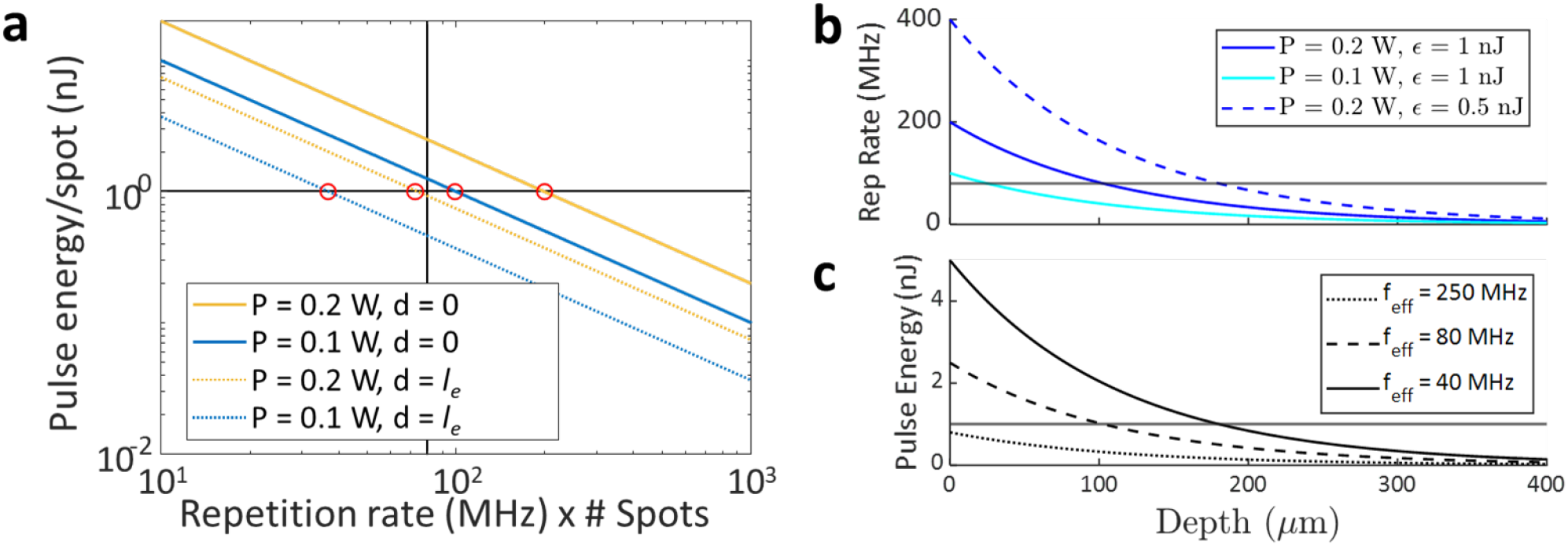
Optimal temporal and spatial splitting. A) At fixed total power, per-spot pulse energy is inversely proportional to effective repetition rate, *f*_eff_ = *f* × *N*_spots_. SNR is increased by increasing focal pulse energy up to the threshold (1 nJ horizontal line shown). Therefore, the optimal effective repetition rate lies at the intersection of the iso-power line with the threshold (circled in red). At a non-zero depth (dotted lines), a lower repetition rate is needed to produce the same focal pulse energy. B) For a single diffraction-limited focal spot, total power of 200 mW, and pulse energy threshold of 1 nJ, the optimal *f*_eff_ goes beneath 80 MHz (horizonal line) at ∼100 μm depth. Decreasing the threshold focal energy (e.g. to 0.5 nJ) increases the optimal repetition rate at a given depth. C) For a fixed *f*_eff_, the focal pulse energy decays exponentially with depth. At *f*_eff_ = 25o MHz, a 1 nJ pulse is not achievable at any depth. At *f*_eff_ = 40 MHz, the pulse energy crosses the 1 nJ threshold at ∼200 μm depth.

This scaling is consistent with a recent report by Sims *et al*. that decreasing laser repetition rate 320-fold from 80 MHz to 250 kHz increased the number of cells imaged at constant SNR from 1 cell/125 mW to 17 cells/125 mW.^5^ Similarly, increasing spot area by a factor of ∼100 to cover an entire cell implies a 10-fold drop in *N*_cells_ compared to our point-scanning estimates (Fig. 3a), predicting fewer than ∼30 cells measurable with an 80 MHz laser with SNR of 10 at the brain surface. Sims *et al*. found that a whole-cell temporally-focused scanless system performed better with speckled illumination that approximated a point array than with more homogeneous illumination,^5^ supporting the view that sparser excitation increases fluorescence count-rate and hence shot noise-limited SNR.

#### Effect of nonlinear saturation

Maximizing focal intensity is only beneficial up to a point. Saturation of the 2P excitation typically occurs at diffraction-limited pulse energies of ∼1 nJ^14^, and nonlinear local photodamage may occur at similar pulse energies.^52,53^ For a fixed pulse duration and spot size, this limit is independent of *f*_eff_. The optimal *f*_eff_ is therefore the repetition rate at which the focal pulse energy density reaches but does not exceed the saturation or damage limit (Fig 6a).

Simultaneously approaching the 2P saturation energy (∼1 nJ) and the maximum power into the sample (∼200 mW) implies a depth-dependent optimum value of *f*_eff_ (Fig 6b). For example, for an 80 MHz laser illuminating the brain with 200 mW, a single focal spot will exceed the 1 nJ limit at the brain surface. Splitting the focus into three diffraction-limited spots (*f*_eff_ = 240 MHz) allows each spot to remain beneath the 1 nJ limit while using the full 200 mW budget. Meanwhile, at a depth of 5 *l*_*e*_, where *l*_*e*_ is the exponential attenuation length, optimal *f*_eff_ ≈ 1 – 10 MHz.^14^ A 2.5 MHz fiber-based soliton laser gave 26-fold greater average signal than an 80 MHz Ti:Sa laser of equal time-average power at depths up to 680 μm in live mouse brain.^14^ In summary, the *f*_eff_ should be tuned to the power limit, focal energy threshold, and imaging depth for each experiment.

The effective repetition rate must also be high enough to visit each measurement point at least once per measurement cycle. For example, an 800 kHz laser could visit 800 points at a 1 kHz revisit rate (assuming a suitable scanner existed) and would provide 100-fold higher time-average count-rate (and 10-fold higher shot noise-limited SNR) than the same laser power delivered at 80 MHz, assuming a sub-saturation pulse energy. Mechanical and acousto-optical scanners have finite slew rates, which can limit the number of cells measurable within 1 ms. When this limit is below the limit set by optical SNR, then beam splitting may increase the number of measurable cells.

#### Optimizing the polarization

For membrane-localized chromophores, signal can be increased by aligning the excitation polarization with the transition dipole of the chromophore.^54^ For 1P excitation, this effect scales as ⟨*cos*^2^θ⟩, where θ is the angle between the excitation polarization and the transition dipole, and the average is taken over the distribution of molecular orientations. The magnitude of the polarization-dependent effect is characterized by 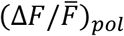, where 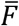 is the fluorescence for unpolarized excitation and 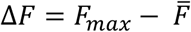, where *F*_*max*_ is the fluorescence for optimal polarization. At the cell periphery, where the optic axis lies in the plane of the membrane, this effect magnitude was 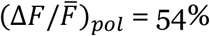 for the dye BeRST1, 20% for ASAP1, 13% for QuasAr3, 12% for ArcLight, and 4.5% for the FRET-opsin GEVI mNeon-Ace.^54^

For 2P excitation, polarization sensitivity scales as ⟨*cos*^4^θ⟩,^55^ and can lead to several-fold polarization-dependent changes in fluorescence from neurites with membrane-bound reporters.^56^ Thus 2P voltage imaging systems could improve their power efficiency substantially by ensuring linearly polarized excitation at the sample and selectively targeting cell membranes that have a favorable orientation relative to the laser polarization; or by modulating polarization during a scan to match the orientation of the target membranes.

### Can advanced analysis techniques overcome the shot noise limit?

Consider the goal of detecting whether a spike occurred (hypothesis *H*^(1)^) or did not occur (*H*^(0)^) during a measurement time *τ*. The mean number of detected photons in the case of a spike is *n*_*1*_, and in the absence of a spike is *n*_0_. The probability distributions for number of detected photons in the two cases are then given by Poisson distributions with means *n*_*1*_ and *n*_*0*_ respectively:

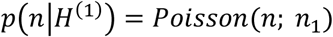

and

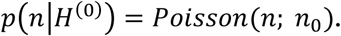

If the number of photons collected in either scenario is not large and the contrast *β* = (*n*_1_ − *n*_0_)/*n*_0_ is also small, then the two probability distributions overlap: a given set of detected photons could have been produced by either the presence or absence of a spike. In such cases, no analysis algorithm can unambiguously determine whether a spike occurred; at best one can determine the relative probabilities of the two hypotheses. This argument is analyzed in detail in Ref. 12.

Voltage signals corresponding to spikes are typically correlated across multiple pixels, and sometimes across frames (depending on the frame rate and spike duration). Since the photon shot-noise is statistically independent between all pairs of pixels, the relative contribution of shot noise can be diminished by weighted averaging across pixels and possibly across frames. If the expected number of photon detections at pixel *i* is ⟨*n*_*i*_⟩ and a filter assigns weight *a*_i_to the pixel, then the expected signal is *S* = Σ_*i*_ *a*_*i*_⟨*n*_*i*_⟩, and the variance in this quantity due to photon shot noise is 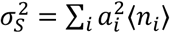. The art in voltage imaging analysis comprises determining the *a*_i_ so as to maximize the difference between *p*(*S*|*H*^(1)^)and *p*(*S*|*H*^(0)^).

Determination of the weights *a*_i_ can be via simple manual or activity-based selections of regions of interest or via optimal detection algorithms, e.g. as in ^57^. When signals from multiple sources overlap, a variety of unmixing algorithms are useful.^58–62^ It is possible even to apply a filter during image acquisition to reduce the data burden.^63^ Recently introduced machine learning algorithms^24,25^ can help determine optimal weighting of pixel signals. None of these techniques, however, overcomes the fundamental uncertainty that different voltages can give identical photon distributions at the detector. Thus claims that these methods “overcome fundamental limits”^23^ of shot noise are misleading.

## Conclusions

Eq. 2 places severe constraints on the number of cells that will be measurable at *d* > 300 μm with 2P voltage imaging, even with substantial improvements in GEVI brightness and voltage sensitivity (Fig. 3c). These findings indicate that 2P imaging of hundreds of neurons with high SNR at depth > 300 μm will require an order-of-magnitude improvement in 2P GEVIs or qualitatively new approaches to imaging. Given the current state of the art, one can maximize SNR and number of measurable cells at depth by using excitation with high numerical aperture, low repetition rate (1 – 10 MHz), short pulses (< 100 fs), optimized polarization, and membrane-targeted illumination with real-time compensation for tissue motion. Optimal imaging can be achieved by customizing spatial multiplexing, repetition rate, and/or excitation NA for the target imaging depth and power limit. Experiments that allow for intermittent imaging and/or distribute the measurements sparsely in space may increase the photothermal limit.

Voltage imaging *in vivo* places stringent demands on the molecular, optical, and data analysis tools. We hope that the above comprehensive analysis of these constraints will shape realistic expectations and guide efforts towards enhancing the performance of 2P voltage imaging.

## Disclosures

The authors have no competing interests to declare.

## Acknowledgements

We thank Andrew Preecha and Shahinoor Begum for technical assistance, and Simon Kheifets, Benjamin Gmeiner, and Vicente Parot for helpful discussions. This work was supported by NIH grant R01-NS126043.

## Code and Data Availability

The instrument control code is available at www.luminosmicroscopy.com and https://github.com/adamcohenlab/luminos-microscopy/.

Data are publicly available on the DANDI archive. Scaling of GEVI fluorescence with 1P and 2P illumination intensity:

https://dandiarchive.org/dandiset/000537.

1P and 2P contrast of JEDI-2P and Voltron2_525_: https://dandiarchive.org/dandiset/000538.

Depth-dependence of fluorescence: https://dandiarchive.org/dandiset/001029.

## Author contributions

HCD, FPB, and AEC designed the study and experiments. HCD collected all of the data. JDW-C performed all mouse surgery and preparation for imaging and assisted with *in vivo* data collection. HCD, and FPB performed all analyses. FPB, HCD, and AEC wrote the manuscript with input from all authors. AEC supervised the work.

## Supporting Information

### Scaling of 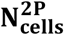 with brightness and voltage sensitivity

Here we derive the dependence of the number of measurable cells on illumination intensity, fluorophore brightness, spike Δ*F*/*F*, measurement bandwidth, and target SNR (Eq. 2 in the main text). We assume a point-scanning 2P illumination system which has perfect targeting to cell membranes, and which can jump between cells with zero delay. This is a best-case scenario: motion artifacts, imperfect targeting, and scanner inertia or finite slew rates further degrade the SNR. We also assume a perfect camera, which converts incident photons to counts with 100% efficiency. We convert camera counts to photons by dividing by the camera quantum efficiency (QE = ∼67% @ 525 nm) and multiplying by the conversion factor (CF = 0.46 photoelectrons/digital count).

Let *F* (counts/s) be the rate of photons collected by a perfect detector from a single cell, let *P* (W) be the laser power at the focus. We define the constant for GEVI brightness, *A*, empirically as the proportionality between squared laser power and fluorescence signal.

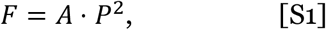

where we have assumed that *P* is changed by adjusting total laser power, keeping laser repetition rate, pulse width, and focal and scan parameters constant.

The number of photons collected during a spike of duration τ is:

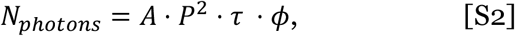

where *ϕ* is the fraction of the time that the laser focus intersects the cell membrane. Let *β* be the fractional change in fluorescence (ΔF/F) during a spike. If there is a contribution to the fluorescence from voltage-insensitive background, then *β* may be smaller than in the background-free case. The signal is:

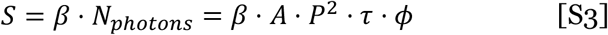

Assuming that the voltage change does not substantially affect the shot noise (i.e. |β| ≪ 1), then the shot noise is:

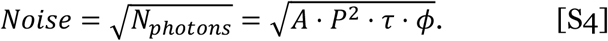

Thus,

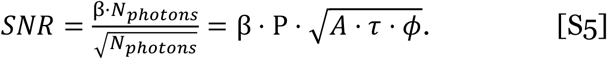

This SNR calculation is applicable only when the laser is targeted to a single cell. If the laser sequentially visits *N* cells, then the duty cycle on each cell is 1/*N*. Assuming zero transit time between cells, the number of photons collected per cell scales inversely with the number of cells:

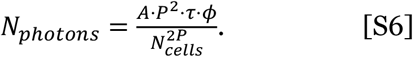

Therefore,

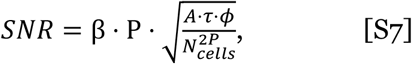

and

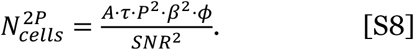

Equation S8 shows the strong dependence of the number of measurable cells on the voltage sensitivity, *β*. The proportionality of SNR and laser power for a single cell holds true for non-scanning excitation modalities as well.^5^

### Scaling of SNR with on and off kinetics

We use the model shown in Fig. 4a. We calculate the endpoint, β, and the area under the rising edge, *R*_*on*_, of the response to a stimulus of length *t* using a reporter with steady-state response, M, and on and off time constants of *τ*_*on*_ and *τ*_*off*_.

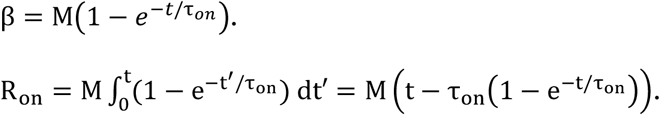

Similarly, we integrate the full area under the decaying response after the voltage pulse. We calculate the integral over the full half-space as the upper bound on response signal.

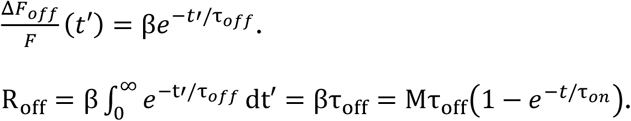

The total response, *R*, is the sum of these two parts:

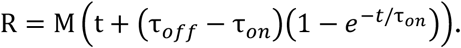

To express SNR, we recognize that *R* takes the place of β ⋅ τ in Eq. S3.

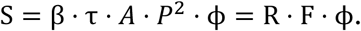

We also set the total integration time, *τ*, in Eq. S4 to be t + τ_*off*_

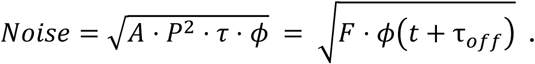

We combine to get:

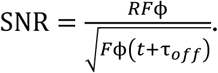

### Theoretical comparison of 1P vs 2P photon efficiencies

We use the properties of JEDI-2P as an exemplary GEVI which works under both 1P and 2P excitation. We assume that the 1P and 2P absorption cross sections of the JEDI-2P chromophore are the same as for eGFP. While this assumption may not be exact, modest variations in these cross sections will not change the conclusion that 2P voltage imaging requires ∼10^4^-fold more power per cell.

First, we estimate the per-molecule excitation rate under 1P excitation. To achieve a per-cell detected digital count rate of 1.5×10^7^ s^-1^ (equivalent to 10^7^ impinging photons/s), required a mean per-cell laser power of 9.6×10^−7^ W, or equivalently an illumination intensity of ∼1 W/cm^2^ (Fig. 1; assuming a HEK cell is approximately 10 μm diameter). This intensity is in the middle of the range used for *in vivo* 1P voltage imaging: recordings of Voltron2 in flies used 200 – 1100 mW/cm^2^,^33^ while high-speed recordings of PV cells in mice used up to 14 W/cm^2^.^33^The decadal molar absorption coefficient of eGFP is ε = 45,000 M^-1^ cm^-1^.^64^ The per-molecule excitation rate is:

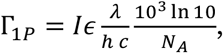

where *I* (W/cm^2^) is the incident intensity, ∈(M^-1^ cm^-1^) is the decadal molar absorption coefficient, *λ* (m) is the wavelength, *h* (6.63×10^−34^ J s) is Planck’s constant, *c* (3×10^8^ m/s) is the speed of light, and *N*_*A*_ (6.02×10^23^ mol^-1^) is Avogadro’s number. We assume *λ* = 488 nm and find that at *I* = 1 W/cm^2^, Γ_1*P*_ = 430 *c*^−1^; at 10 W/cm^2^, Γ_1*P*_ = 4300 *s*^−1^.

Assuming an overall 10% total light collection efficiency (reasonable for a high NA optical system), 10^7^ collected photons/s corresponds to 10^8^ emitted photons/s. At a per-molecule emission rate of 430 s^-1^, this implies 2.3×10^5^ molecules/cell. At a typical HEK cell membrane surface area of 1000 μm^2^,^65^ the density of reporters is 230 μm^-2^.

We now estimate the 2P power needed to match the emitted count rate of 10^8^ s^-1^. The probability that a fluorophore is electronically excited by a single pulse from a 2P optical system is:^66^

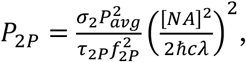

where σ_2_ is the 2P absorption cross section (m^4^ s), *P*_*avg*_ (W) is the time-average power from the laser, *τ*_2*P*_ (s) is the pulse duration, *f*_2 *P*_ is the laser repetition frequency, *NA* is the objective lens numerical aperture, and ℏ, *c*, and λ are as above. The per-molecule rate of excitation is Γ_2 *P*_ = *f*_2*P*_*P*_2*P*_.

We assume parameters typical of a 2P imaging experiment: NA = 1, λ = 920 nm, *τ*_2*P*_ = 200 fs, *f*_2*P*_ = 80 MHz. The 2P absorption cross section of eGFP is σ _*2*_ = 39 GM (39×10^−58^ m^4^ s).^64^

The brightest signal from the cell arises when the laser focus intersects an equatorial membrane, so the optical axis lies in the plane of the membrane, as in Fig. 4c. In this case the membrane area that is optically excited is approximately *A*_2*P*_ = *w*_0_*b*, where *w*_0_ is the waist of the Gaussian focus and *b* is the depth of focus. The focus waist is approximately 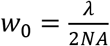, and the depth of focus is 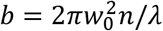, where *n* = 1.33 is the index of refraction. This estimate yields *A*_2P_ ∼ 1 μm^2^, implying that ∼230 reporter molecules are in the 2P focus. To achieve a total emitted photon rate of 10^8^ s^-1^ then implies a per-molecule emission rate of Γ _2P_ = 4.3×10^5^ s^-1^. The time-average laser power to achieve this count rate is 8 mW, 10^4^-fold higher than the 1P power to achieve the same count rate. In the minimal SNR limit of 2.5×10^5^ detected photons/s/cell (corresponding to 2.5×10^6^ emitted photons/s/cell), the minimum time-average 2P power per cell is 0.4 mW (assuming ϕ = 1).

## Biography

### Phil Brooks

studied chemistry at Princeton University, and then completed a PhD in chemistry with Adam Cohen at Harvard University. His work in the Cohen Lab involved developing biological, theoretical, and software tools for robust voltage imaging, and he has abiding interests in tool development for biological research.

### Hunter Davis

received his PhD in chemistry in Mikhail Shapiro’s lab at Caltech. He pursued postdoctoral training with Adam Cohen at Harvard University and is now co-founder and chief scientific officer of a biotech startup, Lorentz Bio.

### David Wong-Campos

was born in Chiclayo, Peru, and attended the Monterrey Institute of Technology and Higher Education in Mexico for his undergraduate. He then completed his Ph. D. in Physics at the University of Maryland under the direction of Christopher Monroe. He took a two-year industry job at a quantum computing startup, IonQ, and held an advisory position at a company, Delee. He returned to academia and is currently pursuing postdoctoral training with Adam Cohen.

### Adam Cohen

is a professor in the Departments of Chemistry and Chemical Biology and Physics at Harvard University. His lab develops physical and chemical tools to study life, with a focus on studying bioelectrical dynamics and the brain. He received PhD degrees from Stanford University and Cambridge University, both in physics, and studied chemistry and physics as an undergraduate at Harvard.

## Notes

### Competing Interest Statement

The authors have declared no competing interest.

### Summary of Updates

More detailed analysis of effects of molecular and instrumentation parameters on performance of 2P voltage imaging.

